# Rumenomics: Evaluation of rumen metabolites from healthy sheep identifies differentially produced metabolites across sex, age, and weight

**DOI:** 10.1101/2025.02.05.636747

**Authors:** Javier Munoz Briones, Brendan K. Ball, Smrutiti Jena, Timothy B. Lescun, Deva D. Chan, Douglas K. Brubaker

## Abstract

**Background:** The rumen harbors a diverse and dynamic microbiome vital in digesting vegetation into metabolic byproducts for energy and general biological function. Although previous studies have reported connections between the rumen and the overall health of the sheep, the exact biological process by which this occurs is not well understood. Therefore, our study aimed to quantify sheep rumen metabolites to determine if enriched biological pathways are differentiable across phenotypic features of sex, age, and weight.

**Results:** We collected and quantified metabolites of rumen samples from sixteen sheep using liquid chromatography-tandem mass spectrometry. We performed a series of univariate and multivariate statistical analyses to interpret the rumen metabolomics data. To identify metabolic pathways associated with the phenotypic features of sex, weight, and age, we used MetaboAnalyst, which identified amino acid metabolism as a distinguishing factor. Among the pathways, phenylalanine metabolism emerged as a key pathway differentiating sheep based on sex and age. Additionally, phenylalanine, tyrosine, and tryptophan biosynthesis were exclusively associated with age. In univariate linear models, we also discovered that these amino acid and protein pathways were associated with weight by age-corrected effect. Finally, we identified arginine and proline biosynthesis as a pathway linked to metabolites with weight.

**Conclusion:** Our study identified differential pathways based on the sex, age, and weight features of sheep. Metabolites produced by the rumen may act as an indicator for sheep health and other ruminants. These findings encourage further investigation of the differentially produced metabolites to assess overall sheep health.

## BACKGROUND

Ruminants like sheep have an evolved digestive system that can break down indigestible plant polysaccharides into usable energy sources. When ruminants consume vegetation, the partially digested plants are anaerobically fermented in the rumen, followed by regurgitation and remastication of the food before the downstream digestive process through the reticulum, omasum, abomasum, and eventually, the small and large intestines. The ability to digest plant cellulose is primarily attributed to the first compartment of the digestive system, referred to as the rumen [1]. The rumen microenvironment harbors a symbiotic relationship between the host and microorganisms such as bacteria, protozoa, and fungi [2].

The host’s rumen microbiome is a multifunctional system that facilitates multiple biological processes. Such pathways include the processing of fibrous feed into metabolic byproducts [3], the production of volatile fatty acids [4], as well as protein and vitamin synthesis [5]. The metabolites produced because of such pathways are essential for physiological functions [6], growth and development [7], and nutrition [8]. The homeostatic function of these pathways ensures the downstream health and wellness of the organism, and if these pathways are dysfunctional, various health conditions may emerge [9]. As a result, understanding how the rumen microbiome influences these pathways will provide an approach to determining a ruminant’s health or condition through the metabolites produced within the rumen.

In efforts to understand the host-microbiome interaction in the rumen, previous studies have used metabolomics to understand the overall health and well-being of ruminants [10–12]. One sheep study reported that metabolites in the rumen microbiota fluctuated at different reproductive periods, potentially affecting health [13]. A survey on grazing lambs found rumen microbial-driven metabolite could potentially regulate downstream body fatty acid metabolism [14]. Another study using Tibetan sheep investigated the relationship between sheep age and metabolite and microbial composition, which showed a potential for microbiome intervention [15].

A challenge in understanding the rumen microbiome could be the complexity of confounding variables and covariates, including but not limited to sex, weight, age, and breed [16,17]. We hypothesized that incorporating covariates to analyze associations of metabolites from the rumen of healthy sheep with their phenotypes would reveal biological and metabolic pathways that inform the relationship between the rumen and produced metabolites. To understand the possible relationship between the rumen microbiome and the metabolites produced, we leveraged multivariate statistical modeling and pathway enrichment analysis to identify potential variables and metabolic pathways that may be identifiable in healthy sheep conditions.

In our study of sheep rumen and their metabolomic profiles, also termed rumenomics, our objective was to determine if the quantified metabolites in rumen varied across healthy sheep populations based on covariates. We paired metabolomics with phenotypic features such as weight, sex, age, and breed to evaluate the functional linkages between 510 metabolites collected from sheep rumen. We performed generalized linear models, partial least squares regression, and multiple hypothesis testing to determine metabolic associations among phenotypic features. With these results, we extracted association insights that revealed the most influential metabolites for each phenotypic feature. This work emphasizes the importance of evaluating livestock well-being, such as sheep, through rumenomics approaches.

## MATERIALS AND METHODS

### Ethics statement

The experimental collection procedures and protocols were conducted on post-euthanasia sheep provided by the Purdue University College of Veterinary Medicine. Sheep were not euthanized for the purpose of this study, but access was granted to allow collection of all samples post-mortem.

### Animals, diets, and experimental design

Sixteen sheep were analyzed in the rumenomics analysis: 10 crossbreeds, 3 Dorset, 2 Hampshire, and 1 Ramboilette. Of these, 9 were male and 7 were female. The Purdue University College of Veterinary Medicine maintained the sheep for 2-4 weeks prior to euthanasia and sample collection in controlled indoor housing conditions as part of a veterinary educational program. Sheep were fed ad libitum hay (75% grass, 25% alfalfa) and had free access to water. The sixteen sheep were healthy, and their age and weight were recorded (**Supplementary Table S1**). On the day of rumen sample collection, sheep were fasted for 24 hours prior to general anesthesia and euthanasia. The sheep were euthanized by intravenous injection of overdose barbiturate (Euthasol, Virbac) while the sheep were under general anesthesia. All pre-euthanasia procedures were performed by the Purdue University College of Veterinary Medicine on an approved, unrelated Institutional Animal Care and Use Committee protocol.

### Rumen fluid sampling and processing

Rumen fluid collection and processing included the use of a portable cooler box, a knife, cheesecloth, 10 L plastic containers, a thermometer, falcon tubes (ranging from 50 to 500 mL), a centrifuge (Eppendorf 5804R), and 1000 mL Kimax media storage glass bottles. Samples of rumen fluid were collected immediately after euthanasia in a necropsy lab to minimize excessive contact between the rumen and air. We filtered the rumen fluid through four consecutive layers of cheesecloth squeezed by hand and transferred into a 10 L plastic container using a funnel. After filtration, 500 mL of rumen fluid was centrifuged in falcon tubes at maximum speed for 10 minutes. After centrifugation, the supernatant was carefully transferred into five 1000 mL Kimax media storage bottles. Rumen fluid samples were divided into two groups: one was stored at – 20 °C for future use. At the same time, the other was prepared for submission to Metabolon (MA, USA) for metabolomics analysis and annotation.

### Sample preparation and metabolomics analysis

Pre-processed rumen fluid samples were centrifuged at 25,000 rpm for 15 minutes, and the supernatant was filtered (0.45 µm filter, Millipore) before being transferred into 2 mL tubes. A total of 48 frozen samples were shipped to Metabolon (MA, USA) for extraction and preparation using their standard solvent extraction methods. The samples were analyzed using Ultrahigh Performance Liquid Chromatography-Tandem Mass Spectrometry (UPLC-MS/MS).

Metabolon’s protocol indicated sample preparation involved the automated MicroLab STAR® system (Hamilton Company). Recovery standards were added to ensure quality control, and proteins were precipitated with methanol under vigorous shaking, followed by centrifugation. The resulting extract was divided into fractions for analysis using different UPLC-MS/MS methods, ensuring coverage of chemically diverse metabolites. These fractions are two for analysis by separate reverse-phase UPLC-MS/MS methods using positive ion mode electrospray ionization, one for analysis by reverse-phase UPLC-MS/MS with negative ion mode electrospray ionization, one for analysis by HILIC/UPLC-MS/MS with negative ion mode electrospray ionization, and one reserved as a backup.

Compounds were identified through Metabolon’s proprietary library, which included retention time/index, the mass-to-charge ratio (m/z), and MS/MS spectral data. Identifications were based on retention index, accurate mass (±10 ppm), and MS/MS scores, allowing precise biochemical differentiation.

### Statistical analysis

We constructed per-metabolite generalized linear models (GLMs) to predict metabolite concentrations based on the phenotypic characteristics of sex, breed, weight, and age, including interaction terms between these phenotypic variables. Metabolites were considered significant if the model p-value was less than 0.05. The false discovery rate (FDR) for model p-values was also calculated for multiple testing corrections. A Wald’s test was applied to assess the relationships between significant metabolites and each phenotypic feature and their interaction terms, with a significance threshold of p-value less than 0.05. Significant metabolites identified by GLMs were further evaluated using the Mann-Whitney test for sex differences and Spearman’s correlation for associations with age, weight, and other significant metabolites. Additionally, linear regression models were constructed to investigate metabolites associated with the interaction between weight and sex in sheep. This multi-faceted approach provided a comprehensive understanding of the metabolite profiles related to the phenotypic traits.

### Multivariate statistical analysis

We implemented a multivariate statistical analysis to study metabolites in rumen fluid using *mixOmics* (R packages) (version 3.19) [18]. Sparse Partial Least Squares Discriminant Analysis (sPLS-DA) was used to determine the metabolite associations with sex, and sparse PLS-regression (sPLS-R) models were created to determine the metabolite associations with age and weight. The model R^2^ and Q^2^, were used to evaluate the validity of the three models to mitigate over-fitting. For the sPLS-DA model, the area under the curve (AUC) indicated the prediction accuracy; higher AUC values closer to 1 reflect higher predictive performance. In the case of sPLS-R, we used the root mean squared error of prediction (RMSEP) to measure model assessment, providing insight into the model’s accuracy in predicting outcomes. In the sPLS models, we calculated variable importance in projection (VIP) to identify important metabolites contributing to the model (VIP > 1). This approach ensured that the pathway enrichment analysis incorporated the most influential metabolites based on their contribution to group separation and variance. Also, we constructed Multiblock PLS models to relate metabolites concentration (X – block) to a three-dimensional Y-block consisting of sheep phenotypic characteristics: sex [female, male], age (months) [mean: 36.81, standard deviation:18.72], and weight [mean: 63.19, standard deviation: 15.19].

### Pathway enrichment analysis with MetaboAnalyst

Metabolites identified as significant from generalized linear models, single sparse PLS, and multiblock-Y PLS were subjected to pathway analysis using MetaboAnalyst (version 6.0) [19]. The pathway analysis module combined results from pathway enrichment analysis with pathway topology analysis. From the 20-pathway library available for mammals, we selected the *Ovis aries* (sheep) for the pathway analysis. Over-representation analysis was performed using a Fisher’s exact test. All statistical p-values from enrichment analysis were further adjusted for multiple testing by Holm-Bonferroni (Holm-adjusted p-value < 0.05) and false discovery rate (FDR-adjusted p-value <L0.05) to correct for multiple hypothesis testing. The impact values indicated the pathway impact values obtained from pathway topology analysis.

## RESULTS

### Rumenomics explored functional linkages between metabolites and sheep phenotypic traits

We collected rumen samples from 16 healthy sheep (9 male and 7 female) immediately postmortem through a veterinary school educational program. We prepared the samples for Metabolon’s metabolomics analysis via liquid chromatography mass spectrometry. Multivariate statistical analyses, including generalized linear models, partial least squares (PLS), and multiblock-Y PLS, were applied to identify if rumen metabolites are coordinately linked to sheep traits such as age, sex, and weight or if they were distinct (**Fig. 1**). The description of sheep phenotypical characteristics are listed in (**Supplementary Table S1**).

**Figure 1.**
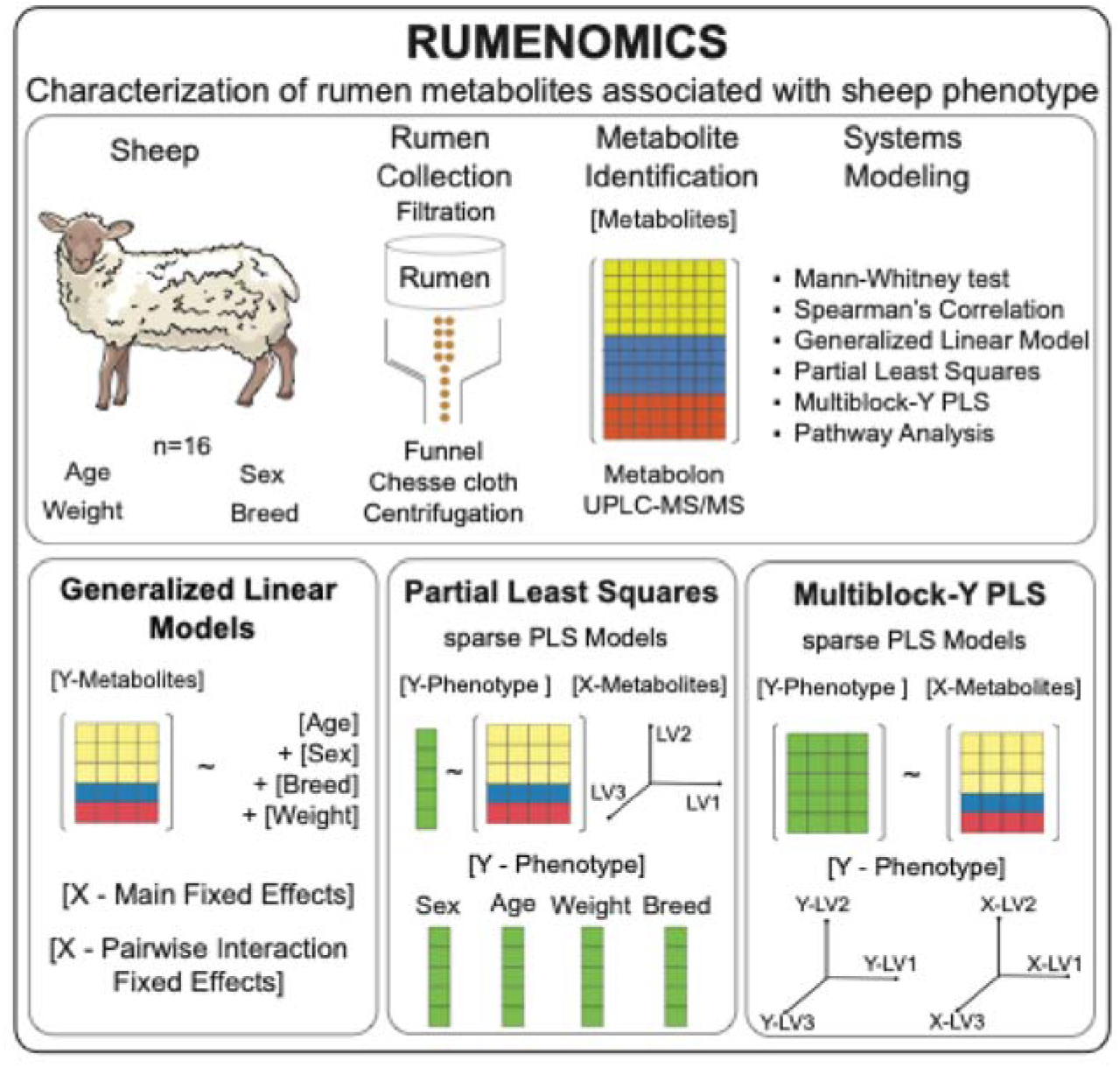
Rumenomics: Evaluation of rumen metabolite content on phenotypic sheep features. Rumen metabolites were collected from 16 healthy sheep, of which 9 were male, and 7 were female. Sheep traits, weight, age, and breed were recorded as covariates. Rumen metabolite content was quantified by ultra-high-performance liquid chromatography-tandem mass spectroscopy by Metabolon. Systems modeling of metabolite content and sheep covariates were determined by PLS and generalized linear model.

### Phenylalanine metabolism was an enriched pathway associated with sheep sex, age, and weight

The main and interaction effect univariate linear models enable the assessment of differentially abundant metabolites by incorporating the effects of sex, breed, weight, and age. These models evaluated individual and combined effects of these phenotypical traits as predictors of metabolite abundance. Based on the p-values from the main effect models, 152 significant metabolites were identified (p-value < 0.05). However, only N-acetylputrescine, gulonate, methionine, and sedoheptulose remained significant after FDR correction for multiple testing (**Fig 2A**). By examining the allocation of these four metabolites within the top 10-ranked significant metabolites (p-value < 0.05) for each phenotypic variable represented in each model coefficient, we identified the following associations: N-acetylputrescine, methionine, and sedoheptulose were significantly associated with sex; N-acetylputrescine and gulonate with weight; and gulonate and sedoheptulose with age (**Supplementary Table S2).**

**Figure 2.**
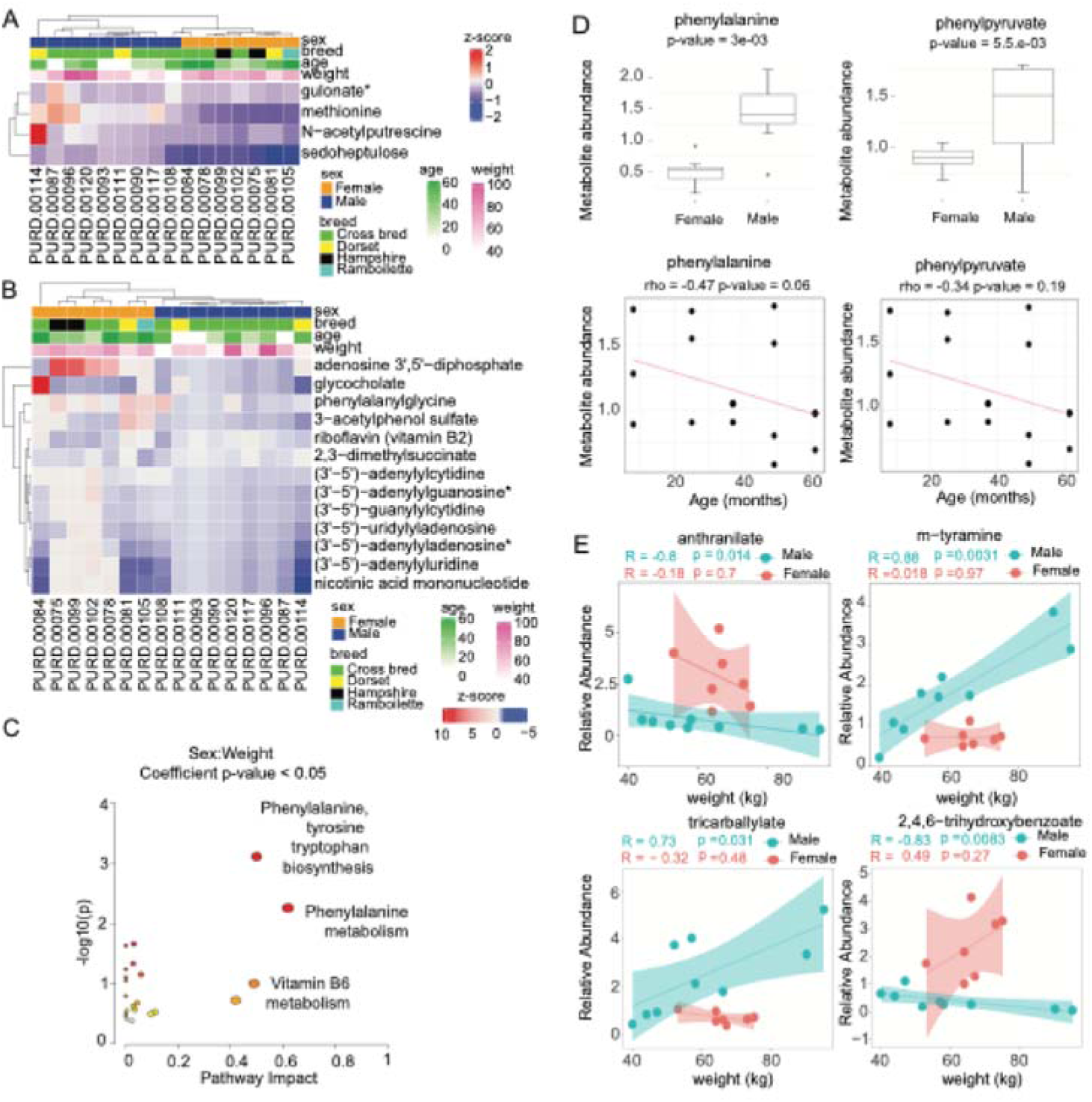
Univariate linear modeling identified metabolites associated with sheep sex, weight, and age. **(A)** Heatmap of significant metabolites (p<0.05) based on main effects univariate linear model. **(B)** Heatmap of significant metabolites (p<0.05) based on age-corrected interaction effects univariate linear model. **(C)** Pathway enrichment analysis performed by MetaboAnalyst using significant metabolites from the interaction term Sex:Weight (p<0.05). (**D**) Mann-Whitney test for sex and Spearman’s correlations with age of phenylalanine and phenylpyruvate. **(E)** Linear regression of metabolites stratified by sheep sex, prioritized by the interaction term Sex:Weight.

Similarly, from the interaction effect models, 90 significant metabolites were identified (p-value < 0.05). After FDR correction, only 17 metabolites remained significant. By examining the allocation of these 17 metabolites across each phenotypic variable represented in each model coefficient, we identified nine significant metabolites associated with sex, weight and age, including: (3’-5’)-adenylyluridine, nicotinic acid mononucleotide (NaMN), 3-acetylphenol sulfate, adenosine 3’,5’-diphosphate, glycocholate, pentadecanoate (15:0), guanosine 5’-diphosphate (GDP), TMP, butyrylputrescine/isobutyrylputrescine (**Fig. 2B**). Exclusive metabolites associated with sex and weight coefficients included: pipecolate, N6-succinyladenosine, N-acetylglycine, 2,4,6-trihydroxybenzoate. Additionally, (3’-5’)-adenylyladenosine and phenethylamine were associated with weight and age, while N-acetylglucosaminylasparagine and sedoheptulose were exclusive with age. The interaction terms Sex:Weight and Sex:Age consistently identified the metabolites that were previously ranked as significant in the interaction effect model (**Supplementary Table S3, Fig. S1**). Interestingly, the interaction term weight:age ranked only five of the 17 metabolites, including N-acetylglucosaminylasparagine, pentadecanoate (15:0), phenethylamine, sedoheptulose and butyrylputrescine/isobutyrylputrescine.

To further study the impact of sex and weight on sheep metabolite profiles both individually and through their interactions, age-corrected linear models were constructed for each metabolite. This approach facilitated the assessment of metabolites associated with sheep weight and sex while treating age as a categorical variable, enabling the detection of non-linear effects across distinct age groups. Based on the p-values from the age-corrected main effect models, 91 significant metabolites were identified (p-value < 0.05). However, no metabolites remained significant after FDR correction for multiple testing. By considering the top-ranked metabolites based on their FDR-adjusted significance, the following top five metabolites were identified from age-corrected main effect models: 5-oxoproline, N-acetylglycine, histidine betaine (hercynine), serotonin, and glycerophosphoglycerol. Metabolites associated with the sex coefficient included N-formylanthranilic acid, ciliatine (2-aminoethylphosphonate), 3-hydroxy-2-ethylpropionate, N,N-dimethyl-5-aminovalerate and methionine. Metabolites related to weight coefficient included histidine betaine (hercynine), serotonin, glycerophosphoglycerol, norvaline, and alpha-hydroxyisovalerate. The complete list of top-ranked metabolites is listed in **Supplementary Table S4.**

Based on the p-values from the age-corrected interaction effect models, 20 significant metabolites were identified (p-value < 0.05). Still, only 13 metabolites remained significant after FDR correction for multiple testing, including, 3-acetylphenol sulfate, nicotinic acid mononucleotide (NaMN), (3’-5’)-uridylyladenosine, (3’-5’)-guanylylcytidine, (3’-5’)-adenylyluridine, (3’-5’)-adenylyladenosine, (3’-5’)-adenylylcytidine, (3’-5’)-adenylylguanosine, adenosine 3’,5’-diphosphate, riboflavin (vitamin B2), glycocholate, 2,3-dimethylsuccinate and phenylalanylglycine. These 13 metabolites were consistently among the top-ranked across the coefficients for sex, weight, and sex:weight interaction. Their relationships with the phenotypic traits of sheep were hierarchically clustered (**Fig. 2B**).

Pathways analysis of metabolites linked to the sex:weight interaction identified ‘phenylalanine, tyrosine and tryptophan biosynthesis’ and ‘phenylalanine metabolism’ as significantly enriched pathways (p-value < 0.05) (**Fig. 2C**). These pathways exhibited the highest impact scores of 0.5 and 0.62, respectively, suggesting that the matched metabolites occupied key structural positions within the pathway network, contributing to their functional influence (**Supplementary Table S5**). On the other hand, significant metabolites associated with the interaction term sex:age from interaction effects univariate linear models identified as significant pathway the ‘pyrimidine metabolism.’ Nevertheless, the pathway impact score was 0.18.

Next, we evaluated how the sex predictor modulated the effect of the weight predictor. A total of 58 metabolites were associated with the sex:weight interaction (p-value < 0.05) (**Supplementary Table S5**). Notably, we found substantial sex-based differences in the abundance of suberate (C8-DC), dodecanedioate (C12), 3-sulfo-alanine, (3’-5’)-adenylyl uridine, (3’-5’)-guanylyl cytidine, 2,3-dimethylsuccinate, anthranilate, 2,4,6-trihydroxybenzoate, riboflavin (vitamin B12), and 4-guanidinobutanoate (p-value < 0.05) (**Fig. S2A)**. Also, Spearman’s correlation analysis revealed significant positive correlations among dinucleotides, including adenosine 3’5’-diphosphate, (3’-5’)-adenylyl guanosine, (3’-5’)-adenyl adenosine, (3’-5’)-adenylyl cytidine, (3’-5’)-adenylyl uridine, (3’-5’)-guanylyl cytidine, and (3’-5’)-uridylyl adenosine (**Fig. S2B**). This finding likely indicated their shared involvement in a specific metabolic pathway related to energy transfer or nucleotide metabolism in the rumen fluid.

Pathways analysis identified phenylalanine and phenylpyruvate as key metabolites for the prioritized pathways. Mann-Whitney analysis showed that these metabolites were significantly more abundant in male sheep. In contrast, Spearman’s correlation analysis revealed a negative correlation with age but no significant correlation with weight (**Fig. 2D**). Subsequently, we constructed linear regression models to examine metabolites associated with the interaction between sex and weight. In male sheep, the relative abundances of m-tyramine, tricarballylate, serotonin, and 3-sulfo-alanine were positively associated with weight. In contrast, anthranilate and 2,4,6-trihydroxybenzoate were negatively associated (**Fig. 2E**). In contrast, no significant associations between metabolite abundances and weight were observed in female sheep.

### Phenylalanine metabolism was linked to sheep sex, whereas phenylalanine, tyrosine, and tryptophan biosynthesis were associated with sheep age

We constructed an sPLS-DA model for sex status and sPLS-R models for age and weight to determine how these covariates influenced changes in metabolite abundance. The scores plot for sPLS-DA sex status model demonstrated clear separation across latent variables (LVs), with LV1 accounting for 37% of the variance in sex and allocating the male sheep in the left side of the score plot (**Fig. 3A**). To identify key metabolites contributing to sex status, we prioritized the top 10 metabolites with the highest Variable Importance in Projection (VIP > 1) scores in the sPLS-DA sex (**Supplementary Table S6A**). The metabolites with the highest VIP score for sex status were sedoheptulose and 3-amino-2-piperidone, which were significantly more abundant in male sheep using the Mann-Whitney test (p-value < 0.05).

**Figure 3.**
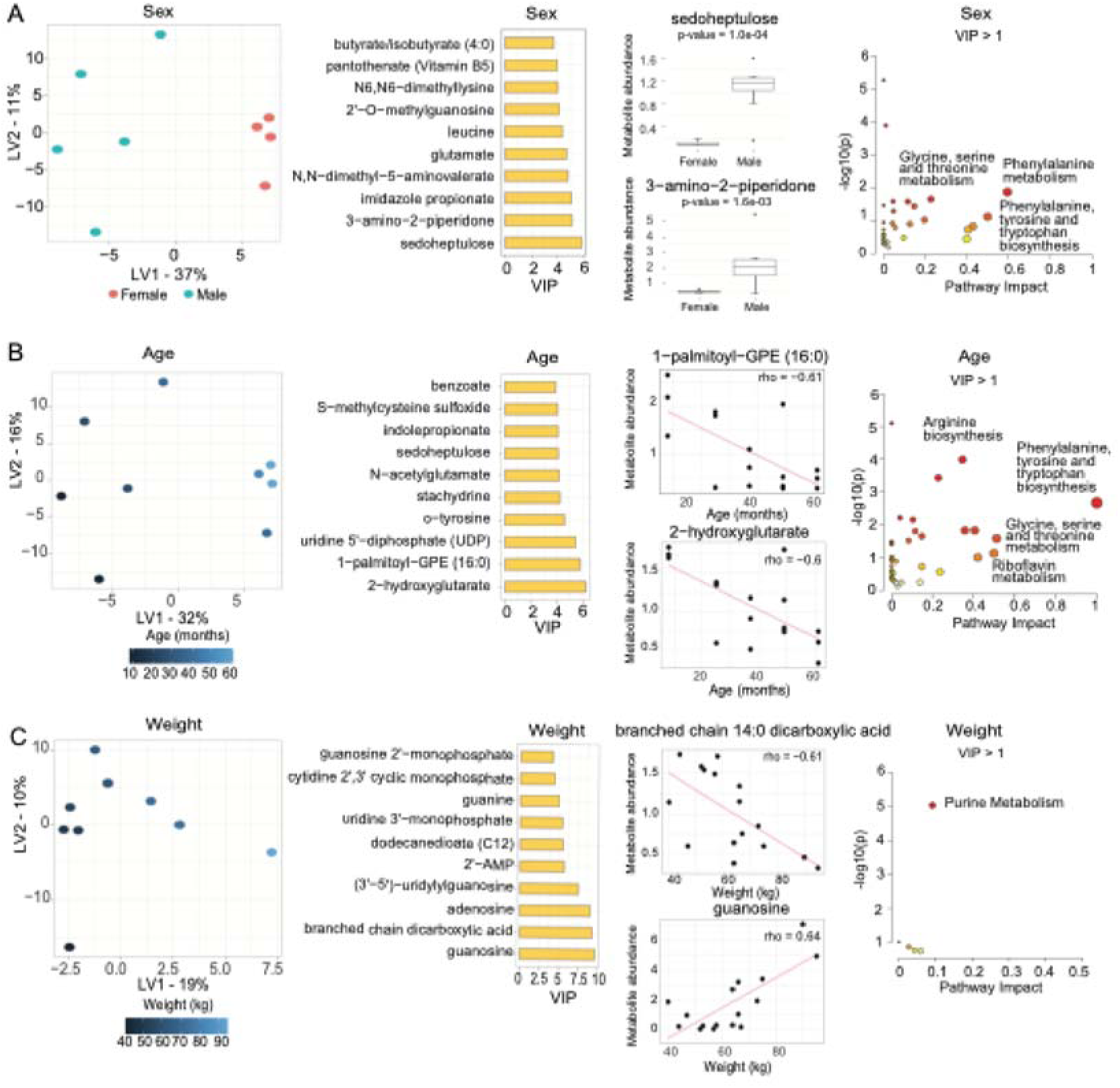
Sparse PLS models were stratified data based on sex, age, and weight. **(A)** Score and VIP plots for sparse PLS by sex. The Mann-Whitney test was applied to the metabolites with the highest VIP score for sex status. Pathway enrichment analysis (MetaboAnalyst) for metabolites with a VIP > 1. **(B)** Score and VIP plots for sparse PLS by sex. Spearman’s correlation analysis was conducted for the metabolites with the highest VIP score for age. Pathway enrichment analysis (MetaboAnalyst) for metabolites with a VIP > 1. **(C)** Score and VIP plots for sparse PLS by weight. Spearman’s correlation analysis was conducted for the metabolites with the highest VIP score for weight. Pathway enrichment analysis (MetaboAnalyst) was performed for metabolites with a VIP > 1.

Pathway enrichment analysis of metabolites with VIP>1 in the sPLS-DA sex model identified ‘phenylalanine metabolism,’ ‘phenylalanine, tyrosine, and tryptophan biosynthesis’, ‘glycine, serine, and threonine metabolism’ as significantly enriched pathways based on the number of metabolites that successfully mapped to these pathways (p-value < 0.05). Among these, ‘phenylalanine metabolism’ had the highest pathway impact score (0.60). However, after multiple hypothesis corrections, only ‘valine, leucine, and isoleucine biosynthesis’ and ‘lysine degradation’ remained significantly enriched (p-value < 0.05). Still, their pathway impact score was negligible (**Fig. 3A**). Based on the loadings of the sPLS sex model and the location of female and male sheep in the score plot, metabolites such as butyrate/isobutyrate, pantothenate, N6, N6-dimethyllysine, 2’-O-methylguanosine, leucine, glutamate, N, N-dimethyl-5-aminovalerate, imidazole propionate, 3-amino-2-piperidone, and sedoheptulose were strongly associated with female sheep, as indicated by their positive loadings in LV1. In contrast, chrysoeriol, 4-chlorobenzoic acid, N-formylanthranilic acid, indole-3-carboxylate, and uridine 5’-diphosphate (UDP) were associated with male sheep due to their negative loadings in LV1 (**Supplementary Table S6B**).

The score plot for the sPLS-R model for age status demonstrated clear separation across latent variables (LVs), with LV1 accounting for 32% of the variance in age (**Fig. 3B**). It allocated the older sheep in the right panel of the score plot. To identify key metabolites contributing to age status, we prioritized the top 10 metabolites with the highest Variable Importance in Projection (VIP > 1) scores in the sPLS-R age (**Supplementary Table S7A**). The metabolites with the highest VIP score for age 2-hydroxyglutarate and 1-palmitoyl-GPR (16:0), both of which were negatively correlated with sheep age using Spearman’s correlation.

Pathway enrichment analysis of metabolites with VIP>1 in the sPLS-R age model identified ‘phenylalanine, tyrosine, and tryptophan biosynthesis’, ‘glycine, serine, and threonine metabolism’, ‘riboflavin metabolism’, and ‘arginine biosynthesis’ as significantly enriched pathways based on the number of metabolites that successfully mapped to these pathways (p-value < 0.05). However, only the pathways’ phenylalanine, tyrosine, and tryptophan biosynthesis’ and ‘arginine biosynthesis’ achieved impact scores of 1 and 0.35, respectively, indicating that the matched metabolites occupy structurally important positions within the pathway network, contributing to their overall structural influence. In contrast, other pathways had impact scores below 0.25 (**Fig. 3B**). Based on the loadings of the sPLS-R age model and the location of old and young sheep in the score plot, metabolites such as 2-hydroxyglutarate, 1-palmitoyl-GPE (16:0), o-tyrosine, stachydrine, N-acetylglutamate, sedoheptulose, S-methylcysteine sulfoxide, benzoate were associated with young sheep given their negative loadings in LV1. In contrast, positive loadings for old sheep were metabolites such as imidazole lactate, indolin-2-one, ophthalmate, phenylalanylanine, 3-hydroxybenzoate, N-formylanthranilic acid, and uridine 5’-diphosphate (UDP) (**Supplementary Table S7B**).

The score plot for sPLS-R model for weight status demonstrated clear separation across latent variables (LVs), with LV1 accounting for 19% of the variance in weight (**Fig. 3C**). The model allocated the sheep with increased weight in the right panel of the score plot. To identify key metabolites contributing to weight status, we prioritized the top 10 metabolites with the highest Variable Importance in Projection (VIP > 1) scores in the sPLS-R weight (**Supplementary Table S8A**). The metabolites with the highest VIP score for weight were guanosine and branched chain (14:0) dicarboxylic acid, in which guanosine was positively correlated with sheep weight. In contrast, using Spearman’s correlation, the branched-chain (14:0) dicarboxylic acid was negatively correlated with sheep weight.

Metaboanalyst with metabolites quantified with a VIP>1 in the sPLS-R weight model identified ‘purine metabolism’ as a significantly enriched pathway (p-value < 0.05); however, its structural influence within the pathway network was negligible (**Fig. 3C**). Based on the loadings of the sPLS-R weight model and the location of sheep with more or less weight in the score plot, the metabolites N-formylphenylalanine, 3-hydroxy-3-methylglutarate, tetradecanedioate (C14), dodecanedioate (C12), and branched-chain 14:0 dicarboxylic acid exhibited negative loading values in LV1, indicating a possible association with sheep with lower weight. Conversely, the metabolites guanosine, adenosine, (3’-5’)-uridylyl guanosine, 2’-AMP, uridine 3’-monophosphate (3’-UMP), guanine, cytidine 2’,3’-cyclic monophosphate, guanosine 2’-monophosphate (2’-GMP), and uridine showed positive loading values in LV1, with a potential connection to sheep of higher weight (**Supplementary Table S8B**).

### Multiblock-Y PLS linked phenylalanine, tyrosine, and tryptophan biosynthesis to sheep sex, age, and weight, and arginine and proline metabolism with sheep weight

We constructed a multiblock-Y PLS model to identify relevant metabolites simultaneously driving separation across sex, age, and weight in healthy sheep. We visualized the score plots for the multiblock-Y PLS model in which LV1 and LV2 with colored labels for sex and age (**Fig. 4A**). The score plots demonstrated a precise classification of sheep sex and age across latent variables LV1, accounting for 50% of the variance among these phenotypes. It allocated the male sheep in the first and fourth quadrants of the score plot and the females in the second and third quadrants.

**Figure 4.**
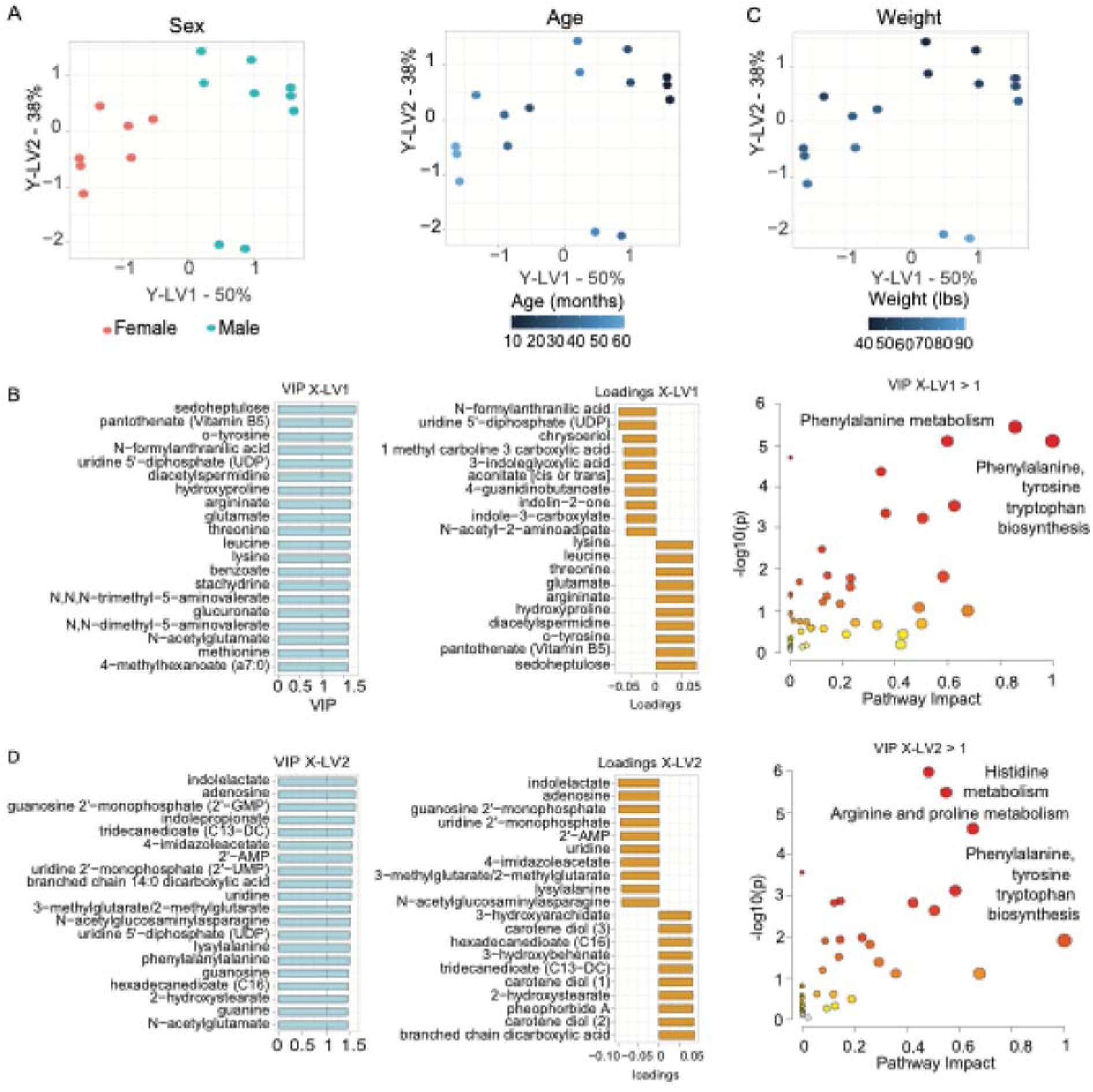
Multiblock-Y PLS model identifying influential metabolites associated with a multivariate response of sex, age, and weight covariates. **(A)** Score plots of Multiblock-Y PLS models by sex and age. **(B)** Top 20 metabolites with the highest VIP scores and the most representative loadings from X-LV1 driving the multivariate PLS decomposition for sex and age, along with pathway analysis using metabolites with VIP scores greater than 1. **(C)** Score plots of Multiblock-Y PLS models by weight. **(D)** Top 20 metabolites with the highest VIP scores and the most representative loadings from X-LV2 driving the multivariate PLS decomposition in weight, along with pathway analysis using metabolites with VIP greater than 1.

To identify key metabolites contributing to the variance in sex and age, we prioritized the top 10 metabolites with the highest Variable Importance in Projection (VIP > 1) scores in multiblock-Y PLS LV1 (**Fig. 4B and Supplementary Table S9A**). The metabolites with the highest VIP score included sedoheptulose, pantothenate (Vitamin B5), o-tyrosine, N-formylanthranilic acid, uridine 5’-diphosphate (UDP), diacetylspermidine, hydroxyproline, argininate, glutamate, threonine.

Interpreting the top representative loadings from multiblock-Y PLS model X-LV1 on the score plots, we identified metabolites with negative loadings, such as N-formylanthranilic acid, uridine 5’-diphosphate, chrysoeriol, 1-methyl-beta-carboline-3-carboxylic acid, 3-indoleglyoxylic acid, aconitate, 4-guanidinolbutanoate, indolin-2-one, indole-3-carboxylate, and N-acetyl-2-aminoaduoate. These metabolites were associated with female sheep in the second and third quadrants on the left side of the plot. In contrast, metabolites with positive loadings, including lysine, leucine, threonine, glutamate, argininate, hydroxyproline, diacetylspermindine, o-tyrosine, pantothenate, and sedoheptulose, influenced the variance towards the male sheep on the right side of the plot, specifically in the first and fourth quadrants. These loadings contributed to the separation for the male sheep (**Fig. 4B**). Interestingly, given that older sheep were located on the left side of the score plot in the second and third quadrants, the metabolites with negative loadings listed above would also be associated with older sheep. Conversely, the metabolites with positive loadings, listed previously, would be related to younger sheep located in the upper right of the first quadrant (**Fig. 4B and Supplementary Table S9B**).

Pathway analysis identified several significantly enriched pathways (p-value < 0.05) based on the number of metabolites that successfully mapped to these pathways, including ‘phenylalanine, tyrosine, and tryptophan biosynthesis’, ‘beta-alanine metabolism’, ‘glycine, serine, and threonine metabolism’, and ‘arginine metabolism’. However, only the pathways ‘phenylalanine tyrosine and tryptophan biosynthesis’, ‘glycine, serine and threonine metabolism’ and ‘arginine biosynthesis’ achieved pathway impact scores of 1, 0.54, and 0.35, respectively, indicating that the matched metabolites occupied structurally essential positions within their respective pathway networks and contributed to their structural influence. While other pathways displayed higher pathway impact scores, they did not have significant matches between the number of metabolites and the pathways (**Fig. 4B and Supplementary Table S10**).

The multiblock-Y PLS modeling revealed that X-LV2 more effectively predicted sheep weight than X-LV1. The first and second quadrants on the upper side of the plot contained sheep with lower weights. Sheep with higher weights were positioned in the third and fourth quadrants at the bottom of the plot (**Fig. 4C**). To identify key metabolites contributing to the variance in weight in LV2, we prioritized the top 10 metabolites with the highest Variable Importance in Projection (VIP > 1) scores in multiblock-Y PLS LV2 (**Supplementary Table S9A**). The metabolites with the highest VIP score included Indolelactate, adenosine, guanosine 2’-monophosphate (2’-GMP), indolepropionate, tridecanedioate (C13-DC), 4-imidazoleacetate, 2’-AMP, uridine 2’-monophosphate (2’-UMP), branched chain 14:0 dicarboxylic acid, uridine (**Fig. 4D**).

Interpreting the top representative loadings from multiblock-Y PLS model X-LV2 on the score plots, we identified metabolites with positive loadings, such as branched chain 14:0 dicarboxylic acid, carotene diol (2), pheophorbide A, 2-hydroxystearate, carotene diol (1), tridecanedioate (C13-DC), 3-hydroxybehenate, hexadecanedioate (C16), carotene diol (3), 3-hydroxyarachidate. These metabolites were associated with sheep with less weight in the first and second quadrants on the upper side of the plot. In contrast, metabolites with negative loadings, including N-acetylglucosaminylasparagine, lysylalanine, 3-methylglutarate/2-methylglutarate, 4-imidazoleacetate, uridine, 2’-AMP, uridine 2’-monophosphate (2’-UMP), guanosine 2’-monophosphate (2’-GMP), adenosine, indolelactate, influenced the variance towards the sheep with more weight on the bottom side of the plot, specifically in the third and fourth quadrants (**Fig. 4D and Supplementary Table S9B**).

Pathway analysis identified several significantly enriched pathways (p-value < 0.05) based on the number of metabolites that successfully mapped to these pathways, including ‘arginine biosynthesis’, ‘histidine metabolism’, ‘arginine and proline metabolism’, and ‘glycine, serine, and threonine metabolism’. These pathways achieved the highest pathway impact scores of 0.45, 0.55, 0.65, and 0.58, respectively, indicating that the matched metabolites occupy structurally essential positions within their respective pathway networks and contribute to their structural influence. Other pathways showed higher pathway impact scores but did not have significant matches between the number of metabolites and the pathways (**Fig. 4D and Supplementary Table S10**).

### Multivariate modeling identified shared metabolic pathways across sex, weight, and age

Identifying common metabolites across generalized linear models, single sPLS models, and multiblock-Y PLS for each phenotypic covariate highlights the consistency of these metabolites’ associations with the sex, age, and weight in sheep. Conversely, the absence of commonalities may indicate a lack of association with these covariates. To further investigate the associations of metabolites with phenotypic covariates revealed by multivariate analysis, we selected enriched pathways with pathway impact scores > 0.5 and p-value < 0.05 for each method associated with a specific phenotype. For instance, in the context of the sheep sex phenotype, we identified pathways top-ranked by the sPLS-DA sex model, the main and interaction effects univariate linear models for sex coefficient, and the age-corrected main and interaction effects univariate linear models for the sex coefficient.

When examining associations between sex and other phenotypic covariates, such as weight and age, we focused on top-ranked pathways enriched from the interaction terms sex:weight and sex:age. These pathways were identified using main and interaction effects univariate linear models, age-corrected main and interaction effects univariate linear models, and the structural insights from Multiblock-Y PLS for LV1, which captured variance related to sex and age. A similar approach was applied to sheep age and weight and their respective interactions. Pathways enriched for these covariates were derived from their respective age, weight coefficients, and interaction terms in univariate linear models. Additionally, we incorporated findings from the sPLS-R age sPLS-R weight models, along with Multiblock-Y PLS results from LV1 for age and LV2 for weight, to gain a comprehensive understanding of these associations. We identified metabolites from enriched pathways associated with sheep sex, age, and weight and their corresponding interactions (**Fig. 5A and Supplementary Table S11**).

**Figure 5.**
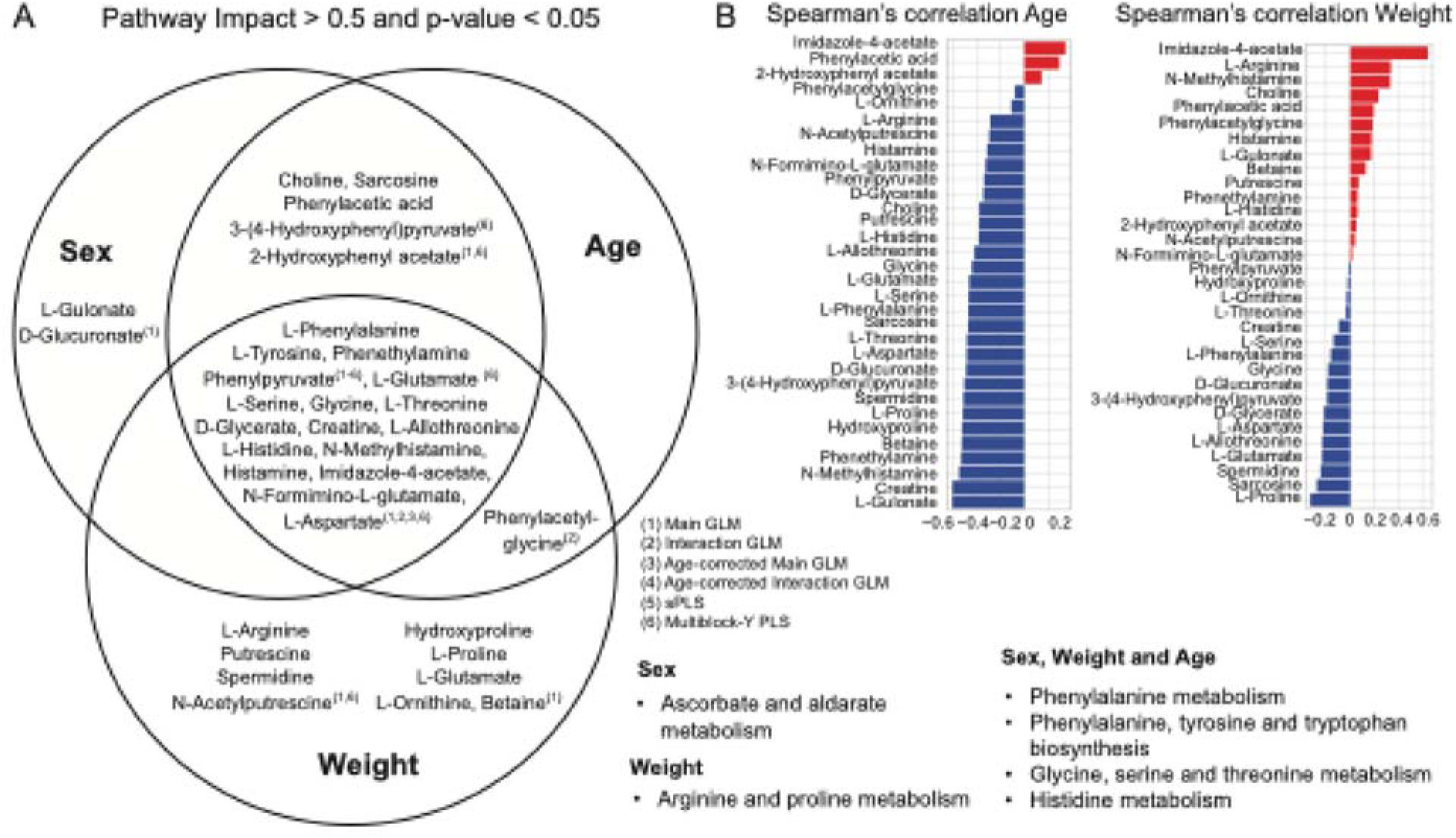
Shared metabolites across sex, weight, and age in multivariate models. **(A)** Venn diagram of metabolites from enriched pathways with an impact score > 0.5 and p-value < 0.05 across analysis, including main GLM, interaction GLM, age-corrected main GLM, age-corrected interaction GLM, sPLS, and Multoblock-Y PLS. **(B)** Spearman’s correlation analysis of common metabolites with sheep age and weight covariate.

Metabolites such as L-phenylalanine, L-tyrosine, phenelthylamine, and phenylpyruvate were consistently reported across multivariate models for sheep sex, age, and weight. These were key metabolites involved in ‘phenylalanine metabolism’, ‘phenylalanine, tyrosine, and tryptophan biosynthesis’. The remaining metabolites in the center of the Venn diagram corresponded to pathways such as ‘glycine, serine and threonine metabolism’ and ‘histidine metabolism.’ Evidence from these associations was obtained from generalized linear models, sPLS, and Multiblock-Y PLS (**Supplementary Table S11**). Similarly, the sex coefficient from the univariate linear model identified L-gulonate and D-glucoronate as key metabolites associated with ‘ascorbate and aldarate metabolism.’ The weight coefficient from the univariate linear model and multiblock-Y PLS identified metabolites listed in the weight-exclusively section of the Venn diagram as being associated with the weight phenotype in sheep, corresponding to ‘arginine and proline metabolism.’ We performed Spearman’s correlation analysis of the top-ranked metabolites from the Venn diagram against weight and age. We found that imidazole-4-acetate showed a strong positive correlation with weight (rho = 0.64) and a moderate positive correlation with age (rho = 0.34). Additionally, creatine and L-gulonate exhibited a strong negative correlation with age (rho = -0.6 for both), while L-gulonate showed a moderate positive correlation with weight (rho = 0.17) and creatine showed a weak negative correlation with weight (rho = -0.1) (**Fig. 5B and Supplementary Table S12**).

## DISCUSSION

In this rumenomics work, we analyzed rumen samples from healthy male and female sheep to identify potential metabolites exhibiting trends across sexes, ages, and weights. We employed various computational modeling approaches, including generalized linear models, partial least squares, Multiblock-Y PLS, Spearman’s correlation analysis, and Mann-Whitney analysis, to identify metabolites differentially present across healthy male and female sheep of varying ages and weights. To interpret the top-ranked metabolites for each sheep phenotype, derived from the coefficients of the generalized linear models, as well as the latent variables distinguishing patterns related to phenotypical features of sex, age, and weight, we used MetaboAnalyst to reveal enriched biological pathways. These pathways were primarily associated with protein and amino acid metabolism among the sheep phenotypic features.

Phenylalanine metabolism emerged as a consistent metabolic pathway associated with sheep sex across multivariate analysis. This pathway is a precursor for numerous metabolites and cannot be synthesized by ruminants and other mammals, requiring the dietary intake of plants and assistance from anaerobic bacteria in the rumen for conversion [20]. The sex-linked differences in phenylalanine metabolism, revealed by multivariate analysis, may be related to variations in rumen microbial communities and hormonal influences, leading to differences in how males and females use this amino acid. Phenylalanine metabolism may support reproductive functions for female ruminants, while in males, it may play a crucial role in growth and muscle development. The sex differences could affect the degradation of phenylalanine into tyrosine or other aromatic metabolites, influencing microbial activity and fermentation efficiency to meet nutritional demands. A study on sheep infused with glucose and insulin demonstrated that insulin-stimulated phenylalanine uptake across the hind limb, indicating that metabolic processes involving phenylalanine are responsive to hormonal changes [21]. These findings suggest opportunities for further research to identify specific microbial species or consortia involved in sex-specific phenylalanine metabolism, explore the impact of phenylalanine supplementation on productivity and health in male and female ruminants, and determine how phenylalanine metabolism interacts with hormonal cycles and reproductive stages in females.

Our computational analysis revealed that the biosynthesis of phenylalanine, tyrosine, and tryptophan and phenylalanine metabolism were strongly associated with age and is likely closely linked to the development of the rumen and its associated functional microbiota composition, which is essential for the fermentation process. At birth, lambs have an undeveloped rumen, but the introduction of solid feed post-weaning promotes microbial colonization [22]. This process facilitates the maturation of the rumen, enabling efficient fermentation and nutrient absorption in young and adult sheep [23]. In this stage, microbial communities can undergo the shikimate pathways to produce these aromatic amino acids [24]. The increased activity in this pathway aligns with the higher protein synthesis demands during growth phases. A study of Tibetan sheep found that phenylethylamine, a metabolite involved in phenylalanine metabolism, was more abundant in sub-adult sheep than in lambs [15]. This suggests that phenylalanine metabolism becomes more pronounced as sheep mature, potentially reflecting the maturation of the rumen and its microbial communities [25,26]. Strikingly, we found mouse and human studies identifying phenylalanine metabolism associated with aging. In a mouse study, phenylalanine plasma increased in mouse models through age [27]. A separate observational study also found serum phenylalanine concentration in humans associated with accelerated telomere shortening, which is related to age-related outcomes [28]. Thus, the biosynthesis of phenylalanine, tyrosine, tryptophan, and phenylalanine metabolism represents key age-related amino acid metabolic pathways.

Arginine and proline metabolism were identified as key metabolic pathways through multiblock-Y, and the main effects were univariate linear models, showing an association with weight. This pathway is central to nitrogen metabolism, influencing ruminants’ nutrient use and overall health [29]. Our findings align with the literature, which suggests that dietary supplementation of arginine (0.25 g arginine for 100g dietary dry feed) improves various bodily functions, including synthesis of proteins, milk production, reproductive performance, growth, and support for metabolic functions [29,30]. Furthermore, a separate group reported that sheep treated with L-arginine via intravenous injection had improved fetal growth and development, highlighting the critical role of arginine in sheep physiology [31].

Purine metabolism was also a pathway identified by sheep weight differences from sPLS modeling. A previous study identified the purine derivatives to creatinine index as a reliable estimate to determine the dry matter intake of livestock [32]. A metabolite identified by univariate linear modeling was riboflavin, also known as vitamin B2. A dietary study on Hu lambs found that the supplementation of riboflavin improved development, digestion, and rumen fermentation [33]. In a mouse study, Tsushima *et al.* reported enhanced purine catabolism in adipose tissue of obese mice [34]. Histidine metabolism is associated with essential amino acids and was upregulated under nutritional diet restriction [35]. In a separate study, free-grazing sheep had elevated L-histidine levels in plasma compared to barn-confined sheep, linking body weight to grazing activity [36]. Thus, further investigation of these metabolic pathways may differentiate phenotypes or health conditions based on anticipated metabolite levels in the rumen accounted for sex, age, and weight.

The Sex:Weight interaction term from the age-corrected univariate linear modeling also identified phenylalanine, tyrosine, and tryptophan biosynthesis, along with phenylalanine metabolism, as metabolic pathways associated with sex and weight. As expected, these pathways play a crucial role in protein synthesis and overall growth in sheep. Increasing dietary intake led to an increase in whole-body protein synthesis, including phenylalanine metabolism [37]. Therefore, weight closely linked to these pathways. Our analysis also revealed sex-specific associations in addition to metabolites and sheep weight, highlighting the complexity of metabolic adaptation in sheep. In male sheep, the relative abundances of m-tyramine, tricarballylate, serotonin, and 3-sulfo-alanine positively associated with weight. M-Tyramine, a trace amine derived from tyrosine, may regulate physiological processes such as neurotransmitter release and vascular tone, which are crucial for growth and metabolism [38]. The increased abundance of tricarballylate, an intermediate in the tricarboxylic acid cycle, could indicate altered energy metabolism during weight gain, as tricarballylate is known to inhibit aconitase, potentially impacting the efficiency of the tricarboxylic acid cycle [39]. Similarly, elevated serotonin levels in heavier males may reflect changes in appetite regulation and gastrointestinal motility, closely tied to body mass [40]. Although the role of 3-sulfo-alanine in metabolism is not well-characterized, its positive association with weight warrants further study.

In contrast, we observed negative associations between weight and the abundances of anthranilate, a tryptophan degradation product [41], and 2,4,6-trihydroxybenzoate, a phenolic compound derivative that may influence antioxidant status and microbial interactions in the rumen [42,43]. The implications of these findings in the context of rumen metabolism need to be further elucidated. Interestingly, no significant associations were detected between metabolite abundance and weight in female sheep, underscoring potential sex-specific metabolic pathways influenced by hormonal and physiological differences. These results suggest that hormonal regulation and body composition differences contribute to sex-specific metabolic responses. Further research is needed to explore these associations and their implications for optimizing sheep health and productivity.

Additionally, sex-based differences in the production of metabolites were identified, with tyrosine being significantly more abundant in male sheep than in females. This is particularly interesting because a reproductive sheep study found that oral administration of L-tyrosine may result in the earlier onset of puberty in male sheep [44]. In a targeted metabolomics study that compared profiles of castrated sheep to control sheep, they also reported L-tyrosine to be significantly less in castrated groups than in control [45]. Furthermore, sedoheptulose 7-phosphate was elevated considerably in control compared to castrated groups [45]. The pathways and metabolites we identified may differentiate across sheep sex in ways that could influence growth, reproductive health, and metabolic efficiency.

While our findings are promising, there are limitations to this study. First, our study looked upon a limited sample size of sheep during a singular point in time, which did not account for longitudinal changes in the rumen microenvironment. The microenvironment within the rumen is dynamic, and thus, other confounding variables may contribute to variations in the rumen [46]. Additionally, we only investigated healthy sheep conditions and did not investigate the potential effects of other health conditions on the abundance of metabolites in the rumen. Thus, further investigation of rumen metabolites of disease or unhealthy conditions may be beneficial in comparing potential changes in the rumen. Addressing these limitations may improve our understanding of the role of metabolites in the rumen produced by ruminants.

## CONCLUSION

The rumen is essential for ruminants like sheep to digest plants for nutrients. Our study examined the metabolites produced in the rumen of healthy sheep to determine if phenotypical features such as sex, age, and weight could alter such quantities. Our computational approach identified significant pathways contributed by differentially produced metabolites across these phenotypes. Understanding the relationship between the host microbiome and the production of metabolites may further inform how different phenotypic features such as sex, age, and weight contribute to the sheep rumen’s dynamic nature and overall health.

## LIST OF ABBREVIATIONS

AUC: Area under the curve
FDR: False discovery rate
PCA: Principal component analysis
PCs: Principal components
PLS: Partial least squares
RMSEP: Root mean squared error of prediction
sPLS: Sparse partial least squares
sPLS-DA: Sparse partial least squares discriminant analysis
sPLS-R: Sparse partial least squares regression
UPLC-MS/MS: Tandem mass spectrometry
VIP: Variance importance in projection

## DECLARATIONS

### Ethics approval

The experimental collection procedures and protocols were conducted on sheep post-euthanasia provided by the Purdue University College of Veterinary Medicine. The use of the sheep prior to euthanasia for educational purposes was approved under an IACUC protocol and was unrelated to the completion of this study.

### Consent for publication

Not applicable for this study.

### Availability of data and material

Generated data for the study is made available in the supplementary files. All code used for the analysis is publicly available at https://github.com/Brubaker-Lab/Rumenomics. The rumenomics data is publicly available at the NIH Common Fund’s National Metabolomics Data Repository, the Metabolomics Workbench, with Study ID ST002908. The data is accessible via its Project DOI: http://dx.doi.org/10.21228/M80X4Q.

### Author’s contributions

JMB, DDC, and DKB conceptualized the study. JMB, BKB, SJ, TBL, DDC, and DKB designed the work. JMB, BKB, SJ, and TBL performed data acquisition. JMB analyzed and visualized the data. JMB and BKB interpreted the data. JMB created the software and code. JMB and BKB wrote the original draft. All authors have read, edited the manuscript, and approved the submitted version.

## Acknowledgments

The authors thank the veterinary students and staff from the Purdue University College of Veterinary Medicine for their support during the collection of rumen samples from the sheep.

---

**Figure S1.**
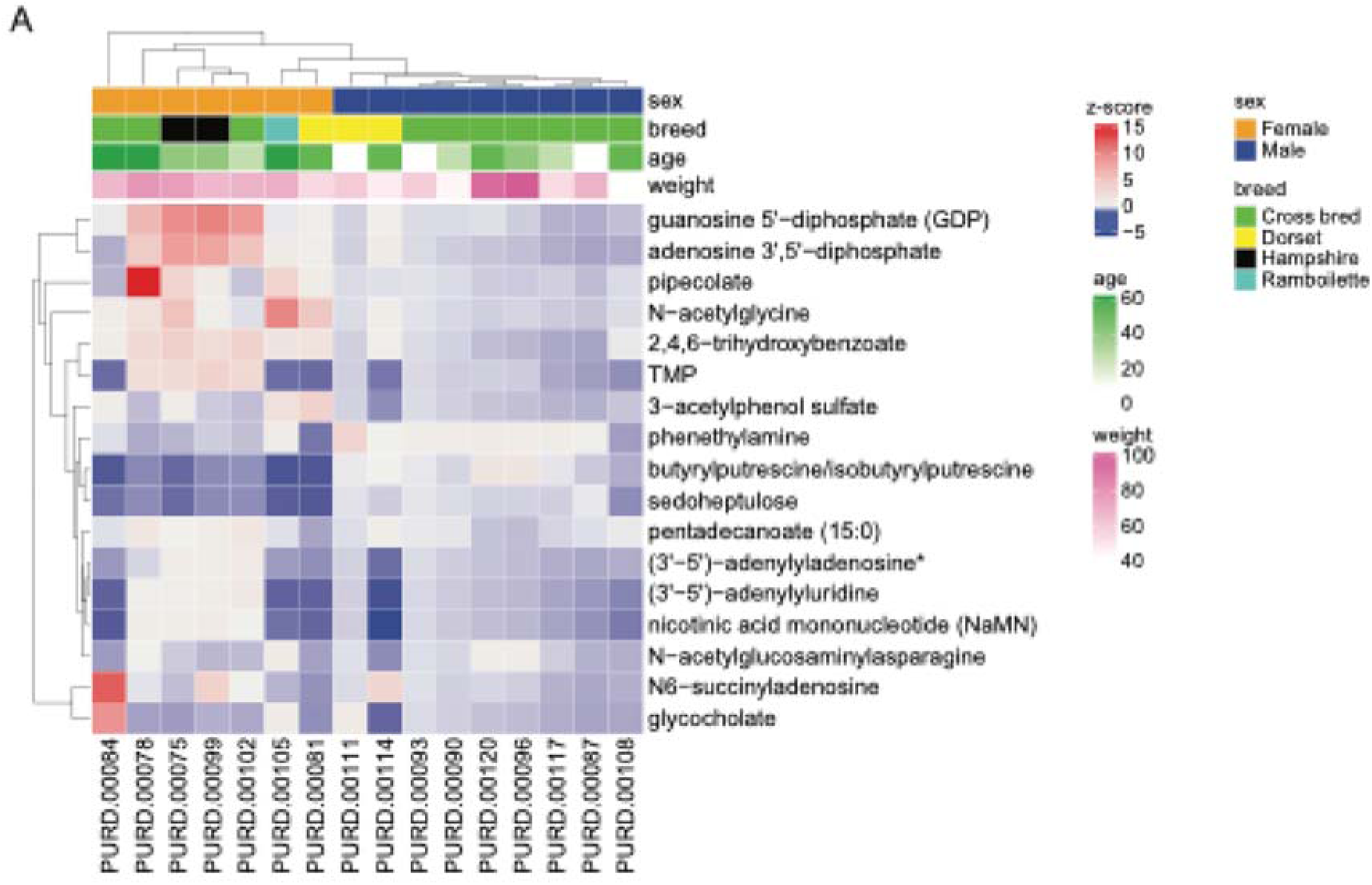
Heatmap of significant metabolites (p<0.05) based on univariate linear modeling identified metabolites associated with sheep sex, weight, and age.

**Figure S2.**
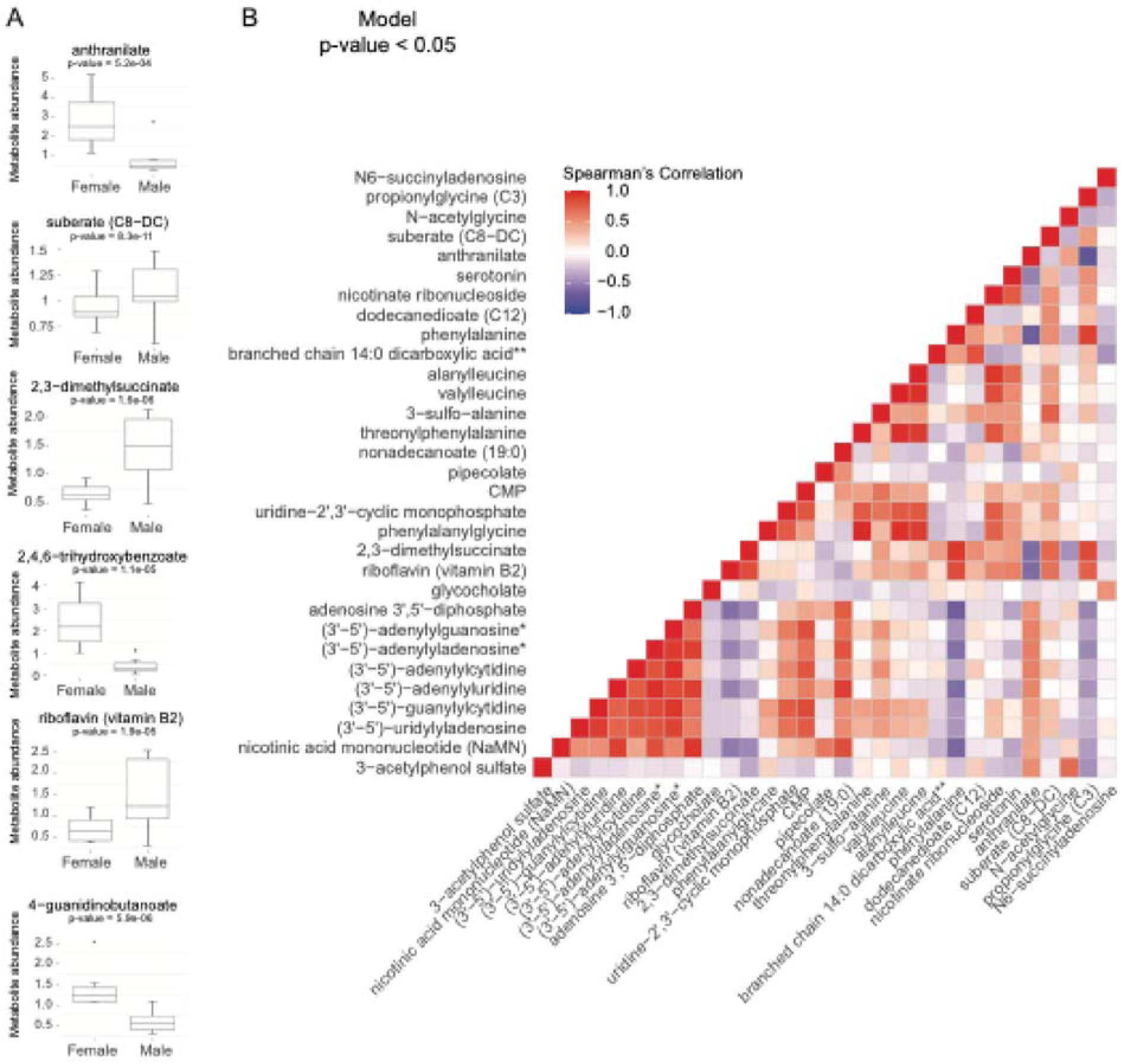
Prioritized metabolites from interaction effects univariate linear modeling. (A) Boxplots of sheep sex-based differences of anthranilate, suberate (C8-DC), 2,3-dimethylsuccinate, 2,4,6-trihydroxybenzoate, riboflavin (vitamin B12), and 4-guanidinobutanoate (p-value < 0.05). (B) Spearman’s correlation analysis from metabolites with model p-values < 0.05.

## REFERENCES

1. Pérez-Barbería FJ. The Ruminant: Life History and Digestive Physiology of a Symbiotic Animal. In: García-Yuste S, editor. Sustainable and Environmentally Friendly Dairy Farms [Internet]. Cham: Springer International Publishing; 2020 [cited 2024 Dec 27]. p. 19–45. Available from: 10.1007/978-3-030-46060-0_2

2. Cholewińska P, Czyż K, Nowakowski P, Wyrostek A. The microbiome of the digestive system of ruminants – a review. Animal Health Research Reviews. 2020;21:3–14.

3. Zhang YK, Zhang XX, Li FD, Li C, Li GZ, Zhang DY, et al. Characterization of the rumen microbiota and its relationship with residual feed intake in sheep. Animal. 2021;15:100161.

4. Cheng J, Zhang X, Xu D, Zhang D, Zhang Y, Song Q, et al. Relationship between rumen microbial differences and traits among Hu sheep, Tan sheep, and Dorper sheep. Journal of Animal Science. 2022;100:skac261.

5. Hackmann TJ, Firkins JL. Maximizing efficiency of rumen microbial protein production. Front Microbiol [Internet]. 2015 [cited 2024 Dec 27];6. Available from: https://www.frontiersin.org/journals/microbiology/articles/10.3389/fmicb.2015.00465/full

6. Moraïs S, Mizrahi I. The Road Not Taken: The Rumen Microbiome, Functional Groups, and Community States. Trends in Microbiology. 2019;27:538–49.

7. Lane MA, Baldwin RL VI, Jesse BW. Sheep rumen metabolic development in response to age and dietary treatments3. Journal of Animal Science. 2000;78:1990–6.

8. Liu K, Zhang Y, Yu Z, Xu Q, Zheng N, Zhao S, et al. Ruminal microbiota–host interaction and its effect on nutrient metabolism. Animal Nutrition. 2021;7:49–55.

9. Magrin L, Brscic M, Lora I, Prevedello P, Contiero B, Cozzi G, et al. Assessment of Rumen Mucosa, Lung, and Liver Lesions at Slaughter as Benchmarking Tool for the Improvement of Finishing Beef Cattle Health and Welfare. Front Vet Sci [Internet]. 2021 [cited 2024 Dec 27];7. Available from: https://www.frontiersin.org/journals/veterinary-science/articles/10.3389/fvets.2020.622837/full

10. Tian H, Wang W, Zheng N, Cheng J, Li S, Zhang Y, et al. Identification of diagnostic biomarkers and metabolic pathway shifts of heat-stressed lactating dairy cows. Journal of Proteomics. 2015;125:17–28.

11. Palma M, Scanlon T, Kilminster T, Milton J, Oldham C, Greeff J, et al. The hepatic and skeletal muscle ovine metabolomes as affected by weight loss: a study in three sheep breeds using NMR-metabolomics. Sci Rep. 2016;6:39120.

12. Zhang J, Gao Y, Guo H, Ding Y, Ren W. Comparative metabolome analysis of serum changes in sheep under overgrazing or light grazing conditions. BMC Vet Res. 2019;15:469.

13. Sha Y, Liu X, Li X, Wang Z, Shao P, Jiao T, et al. Succession of rumen microbiota and metabolites across different reproductive periods in different sheep breeds and their impact on the growth and development of offspring lambs. Microbiome. 2024;12:172.

14. Li Z, Zhao X, Jian L, Wang B, Luo H. Rumen microbial-driven metabolite from grazing lambs potentially regulates body fatty acid metabolism by lipid-related genes in liver. Journal of Animal Science and Biotechnology. 2023;14:39.

15. Li H, Yu Q, Li T, Shao L, Su M, Zhou H, et al. Rumen Microbiome and Metabolome of Tibetan Sheep (Ovis aries) Reflect Animal Age and Nutritional Requirement. Front Vet Sci [Internet]. 2020 [cited 2024 Dec 27];7. Available from: https://www.frontiersin.org/journals/veterinary-science/articles/10.3389/fvets.2020.00609/full

16. González-Recio O, Martínez-Álvaro M, Tiezzi F, Saborío-Montero A, Maltecca C, Roehe R. Invited review: Novel methods and perspectives for modulating the rumen microbiome through selective breeding as a means to improve complex traits: Implications for methane emissions in cattle. Livestock Science. 2023;269:105171.

17. Chaucheyras-Durand F, Ossa F. REVIEW: The rumen microbiome: Composition, abundance, diversity, and new investigative tools. The Professional Animal Scientist. 2014;30:1–12.

18. Rohart F, Gautier B, Singh A, Cao K-AL. mixOmics: An R package for ‘omics feature selection and multiple data integration. PLOS Computational Biology. 2017;13:e1005752.

19. Pang Z, Lu Y, Zhou G, Hui F, Xu L, Viau C, et al. MetaboAnalyst 6.0: towards a unified platform for metabolomics data processing, analysis and interpretation. Nucleic Acids Research. 2024;52:W398–406.

20. Allison MJ. Phenylalanine biosynthesis from phenylacetic acid by anaerobic bacteria from the rumen. Biochemical and Biophysical Research Communications. 1965;18:30–5.

21. Wester TJ, Lobley GE, Birnie LM, Lomax MA. Insulin Stimulates Phenylalanine Uptake across the Hind Limb in Fed Lambs. The Journal of Nutrition. 2000;130:608–11.

22. Baldwin RL, McLeod KR, Klotz JL, Heitmann RN. Rumen Development, Intestinal Growth and Hepatic Metabolism In The Pre- and Postweaning Ruminant. Journal of Dairy Science. 2004;87:E55–65.

23. Herath HMGP, Pain SJ, Kenyon PR, Blair HT, Morel PCH. Rumen Development of Artificially-Reared Lambs Exposed to Three Different Rearing Regimens. Animals (Basel). 2021;11:3606.

24. Hassan F, Guo Y, Li M, Tang Z, Peng L, Liang X, et al. Effect of Methionine Supplementation on Rumen Microbiota, Fermentation, and Amino Acid Metabolism in In Vitro Cultures Containing Nitrate. Microorganisms. 2021;9:1717.

25. Xu Y, Feng T, Ding Z, Li L, Li Z, Cui K, et al. Age-related compositional and functional changes in the adult and breastfed buffalo rumen microbiome. Front Microbiol [Internet]. 2024 [cited 2024 Dec 27];15. Available from: https://www.frontiersin.org/journals/microbiology/articles/ 10.3389/fmicb.2024.1342804/full

26. Du S, Bu Z, You S, Jiang Z, Su W, Wang T, et al. Integrated rumen microbiome and serum metabolome analysis responses to feed type that contribution to meat quality in lambs. Animal Microbiome. 2023;5:65.

27. Czibik G, Mezdari Z, Murat Altintas D, Bréhat J, Pini M, d’Humières T, et al. Dysregulated Phenylalanine Catabolism Plays a Key Role in the Trajectory of Cardiac Aging. Circulation. 2021;144:559–74.

28. Eriksson JG, Guzzardi M-A, Iozzo P, Kajantie E, Kautiainen H, Salonen MK. Higher serum phenylalanine concentration is associated with more rapid telomere shortening in men12. The American Journal of Clinical Nutrition. 2017;105:144–50.

29. Wu G, Bazer FW, Satterfield MC, Gilbreath KR, Posey EA, Sun Y. L-Arginine Nutrition and Metabolism in Ruminants. Adv Exp Med Biol. 2022;1354:177–206.

30. Gootwine E, Rosov A, Alon T, Stenhouse C, Halloran KM, Wu G, et al. Effect of supplementation of unprotected or protected arginine to prolific ewes on maternal amino acids profile, lamb survival at birth, and pre- and post-weaning lamb growth. Journal of Animal Science. 2020;98:skaa284.

31. Lassala A, Bazer FW, Cudd TA, Datta S, Keisler DH, Satterfield MC, et al. Parenteral Administration of l-Arginine Enhances Fetal Survival and Growth in Sheep Carrying Multiple Fetuses123. J Nutr. 2011;141:849–55.

32. Del Valle TA, de Morais JPG, Campana M, Azevedo EB, Louvandini H, Abdalla AL. Purine derivatives and creatinine urine excretion as a tool to estimate sheep feed intake. Animal Feed Science and Technology. 2023;301:115666.

33. Ren N, Zhang X, Hao X, Dong Y, Wang X, Zhang J. Effect of Dietary Inclusion of Riboflavin on Growth, Nutrient Digestibility and Ruminal Fermentation in Hu Lambs. Animals (Basel). 2022;13:26.

34. Tsushima Y, Nishizawa H, Tochino Y, Nakatsuji H, Sekimoto R, Nagao H, et al. Uric Acid Secretion from Adipose Tissue and Its Increase in Obesity. J Biol Chem. 2013;288:27138–49.

35. Feng T, Ding H, Wang J, Xu W, Liu Y, Kenéz Á. Metabolite Profile of Sheep Serum With High or Low Average Daily Gain. Front Vet Sci. 2021;8:662536.

36. Wang B, Luo Y, Su R, Yao D, Hou Y, Liu C, et al. Impact of feeding regimens on the composition of gut microbiota and metabolite profiles of plasma and feces from Mongolian sheep. J Microbiol. 2020;58:472–82.

37. Harris PM, Skene PA, Buchan V, Milne E, Calder AG, Anderson SE, et al. Effect of food intake on hind-limb and whole-body protein metabolism in young growing sheep: chronic studies based on arterio-venous techniques. Br J Nutr. 1992;68:389–407.

38. Berry MD. The potential of trace amines and their receptors for treating neurological and psychiatric diseases. Rev Recent Clin Trials. 2007;2:3–19.

39. Russell JB, Forsberg N. Production of tricarballylic acid by rumen microorganisms and its potential toxicity in ruminant tissue metabolism. Br J Nutr. 1986;56:153–62.

40. Yano JM, Yu K, Donaldson GP, Shastri GG, Ann P, Ma L, et al. Indigenous Bacteria from the Gut Microbiota Regulate Host Serotonin Biosynthesis. Cell. 2015;161:264–76.

41. Oxenkrug GF. Tryptophan kynurenine metabolism as a common mediator of genetic and environmental impacts in major depressive disorder: the serotonin hypothesis revisited 40 years later. Isr J Psychiatry Relat Sci. 2010;47:56–63.

42. Van Soest P. Nutritional Ecology of the Ruminant by Peter J. Van Soest | Hardcover [Internet]. 2nd ed. Cornell University Press; 1982 [cited 2024 Dec 27]. Available from: https://www.cornellpress.cornell.edu/book/9780801427725/nutritional-ecology-of-the-ruminant/

43. Sankaranarayanan R, Valiveti CK, Kumar DR, Van Slambrouck S, Kesharwani SS, Seefeldt T, et al. The Flavonoid Metabolite 2,4,6-Trihydroxybenzoic Acid Is a CDK Inhibitor and an Anti-Proliferative Agent: A Potential Role in Cancer Prevention. Cancers (Basel). 2019;11:427.

44. El-Battawy K. Reproductive and Endocrine Characteristics of Delayed Pubertal Ewe-Lambs after Melatonin and l-Tyrosine Administration. Reproduction in Domestic Animals. 2006;41:1– 4.

45. Li J, Tang C, Zhao Q, Yang Y, Li F, Qin Y, et al. Integrated lipidomics and targeted metabolomics analyses reveal changes in flavor precursors in psoas major muscle of castrated lambs. Food Chemistry. 2020;333:127451.

46. Wang L, Zhang K, Zhang C, Feng Y, Zhang X, Wang X, et al. Dynamics and stabilization of the rumen microbiome in yearling Tibetan sheep. Sci Rep. 2019;9:19620.

